# Assessment of Variability in the Plasma 7k SomaScan Proteomics Assay

**DOI:** 10.1101/2022.09.14.507978

**Authors:** Julián Candia, Gulzar N. Daya, Toshiko Tanaka, Luigi Ferrucci, Keenan A. Walker

**Affiliations:** Translational Gerontology Branch, National Institute on Aging, National Institutes of Health, Baltimore, MD 21224, USA; Laboratory of Behavioral Neuroscience, National Institute on Aging, National Institutes of Health, Baltimore, MD 21224, USA

## Abstract

SomaScan is a high-throughput, aptamer-based proteomics assay designed for the simultaneous measurement of thousands of proteins with a broad range of endogenous concentrations. In its most current version, the 7k SomaScan assay v4.1 is capable of measuring 7,288 human proteins. In this work, we present an extensive technical assessment of this platform based on a study of 2,050 samples across 22 plates. Included in the study design were inter-plate technical duplicates from 102 human subjects, which allowed us to characterize different normalization procedures, evaluate assay variability by multiple analytical approaches, present signal-over-background metrics, and discuss potential specificity issues. By providing detailed performance assessments on this wide range of technical aspects, we aim for this work to serve as a valuable resource for the growing community of SomaScan users.

## Introduction

SomaScan^1, 2^ (spelled SOMAscan in earlier versions) is a highly multiplexed, aptamer-based assay capable of simultaneously measuring thousands of human proteins broadly ranging from femto- to micro-molar concentrations. This platform relies upon a new generation of protein-capture SOMAmer (*Slow Offrate Modified Aptamer*) reagents^3^. SOMAmers are based on single-stranded, chemically-modified nucleic acids, selected via the so-called SELEX (*Systematic Evolution of Ligands by EXponential enrichment*) process, which is designed to optimize high affinity, slow off-rate, and high specificity to target proteins. These targets extensively cover major molecular functions including receptors, kinases, growth factors, and hormones, and span a diverse collection of secreted, intracellular, and extracellular proteins or domains. In recent years, SomaScan has increasingly been adopted as a powerful tool for biomarker discovery across a wide range of diseases and conditions, as well as to elucidate their biological underpinnings in proteomics and multi-omics studies^4–11^.

Concurrently with its wider adoption, the proteome coverage of SomaScan has increased over time, from roughly 800 SOMAmers in 2009, to 1,100 in 2012, 1,300 in 2015, 5,000 in 2018, and the most recent 7,000 protein assay available since 2020. Using 2,624 samples analyzed between 2015 and 2017, our teams at the U.S. National Institutes of Health (NIH) performed in-depth analyses to assess the technical features of the 1.1k and 1.3k SomaScan assays, including normalization procedures and their variability^12^, later followed by technical assessments from other laboratories^13–16^. To the best of our knowledge, however, no comprehensive technical assessments have been performed on the newest 7k SomaScan assay.

In this context, the goal of our work is to fill this knowledge gap by performing an extensive analysis of normalization approaches and technical variability across more than 7,000 SOMAmers from the most current version of this assay, thereby providing a valuable update on previous efforts to characterize the assay’s performance. Moreover, since most of the SOMAmers in earlier versions are contained in the newest one (Supplementary Figure 1, Supplementary Data 1), this study will be relevant to investigations based on those earlier assay versions as well. Our work is based on a major proteomics profiling effort from a total of 2,050 plasma samples spanning 22 plates, designed with inter-plate technical duplicates from 102 human subjects. Utilizing this extensive dataset, we describe and evaluate different normalization procedures, assess precision and variability by multiple analytical approaches, determine signal-over-background metrics across the assay, and discuss potential specificity issues. As SomaScan becomes more widely adopted and utilized as a state-of-the-art tool for proteomics discovery, we hope that this work will constitute a cornerstone technical reference and resource for future studies.

## Results

Fig. 1 shows Principal Component Analysis (PCA) of buffer wells, calibrators, QC, and human donor samples distributed across 22 plates, shown in different colors based on plate (see Methods section for details). Panels (a-b) display PCA plots obtained from raw data, whereas (c-d) correspond to fully normalized (hyb.msnCal.ps.cal.msnAll) data. In the left-side panels, PC1 appears dominated by the difference between buffer-only wells and those that contain samples (either control or experimental). Indeed, PC1 captures most of the overall variance (from 64.7% in the raw data to 59.5% in the fully normalized data). In order to observe finer details of variability among samples, the right-side panels show PCA plots after removing the buffer-only wells. We observe that, as a result of normalization, variance is more evenly distributed in Fig. 1(d), with the amount of variance explained by PC1 less than half of that in Fig. 1(b). In Fig. 1(d), we see that PC2 captures the differences between experimental and control (calibrator and QC) samples. These differences, in contrast, are obscured in Fig. 1(b) by nuisance effects that have not been removed from the raw data.

**Figure 1.**
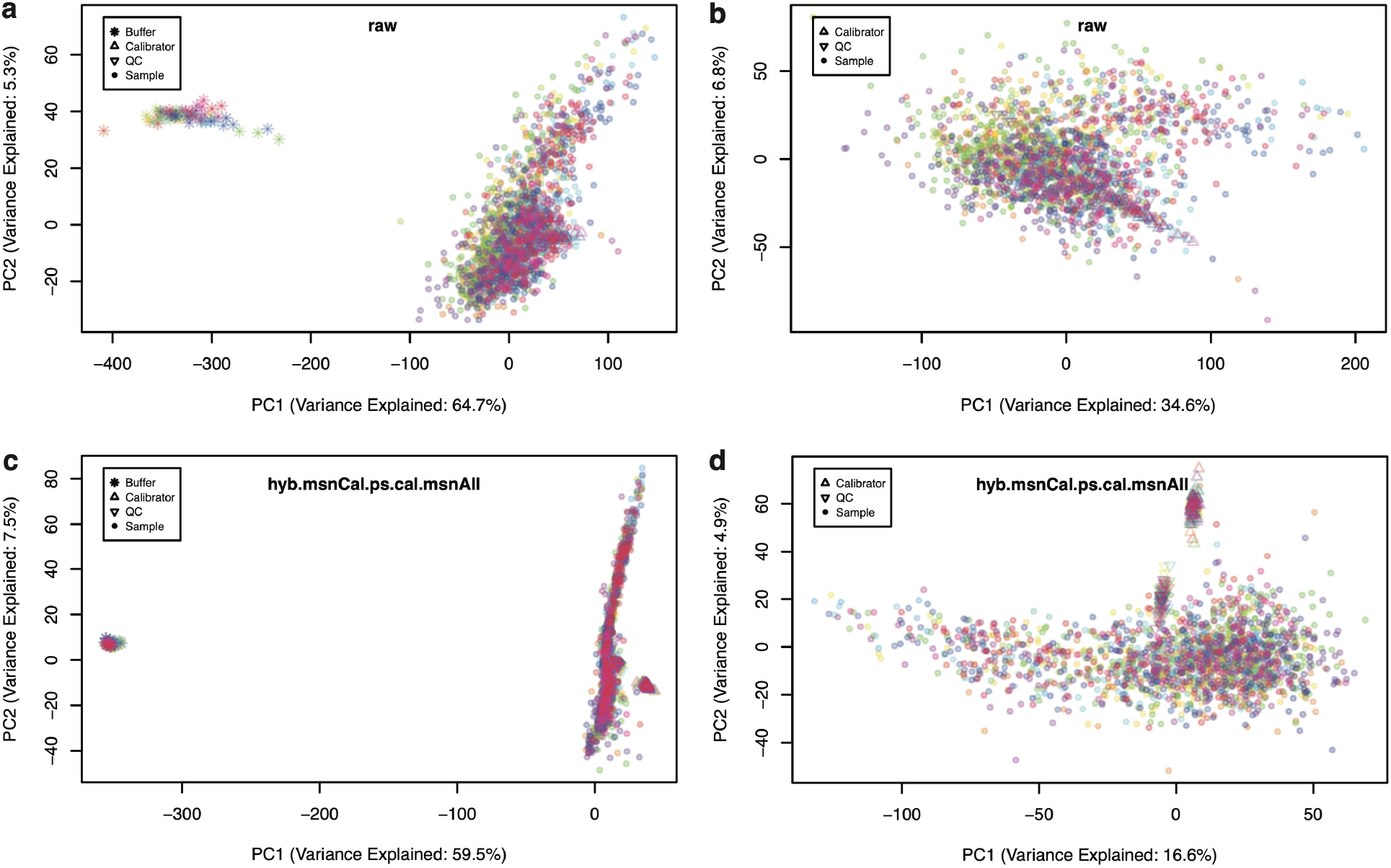
Principal component analysis (PCA) of 2,043 samples. PCA comprising 68 buffers, 110 calibrators, 66 QC and 1,799 human donor samples distributed across 22 plates. Each plate is shown in a different color. **a** Raw data with all sample types. **b** Raw data with buffer wells removed. **c** Fully normalized dataset (hyb.msnCal.ps.cal.msnAll) with all sample types. **d** Fully normalized dataset with buffer wells removed.

Although PCA results in Fig. 1 appear to highlight the importance of normalization to remove unwanted variance, it is possible for normalization procedures to overfit the data, whereby introducing spurious sources of noise in the attempt to remove genuine ones. In order to perform an independent assessment of variability along the normalization process, we shall focus on duplicate biological inter-plate pairs from samples obtained from *n*_*dupl*_ = 102 human participants.

Uncertainties (defined as expected or predicted differences among technical replicates) are typically modeled as combinations of additive terms that are constant throughout the range of measurement (e.g. background noise) and multiplicative factors that scale linearly with the concentration (e.g. volume differences). The metric used to measure variation should be consistent with the uncertainty model assumed. Typically, the overall uncertainty is dominated at low concentrations by additive effects, transitioning at higher concentrations to multiplicative effects. Fig. 2 shows Bland-Altman plots of RFU absolute differences between technical duplicates as a function of their mean, for all duplicate pairs and all human protein SOMAmers, displayed for (a) raw and (b) fully normalized data. The spread increases with relative concentration, suggesting that scaled relative differences, as defined by

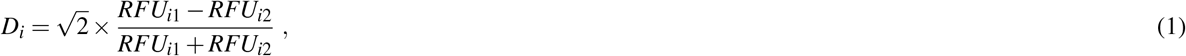

where *i* = 1, …, *n*_*dupl*_, may be used as appropriate building blocks of a variability metric^17^. Notice that scaled relative differences are dimensionless; the 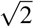 prefactor is added because the differences arise from imprecision in both measurements, which adds quadratically, whereas the desired result is the variability of one measurement. With *n*_*dupl*_ technical duplicate pairs, different variability estimates may be built, as follows. The Root-Mean-Squared Variation (RMSV) metric is defined as

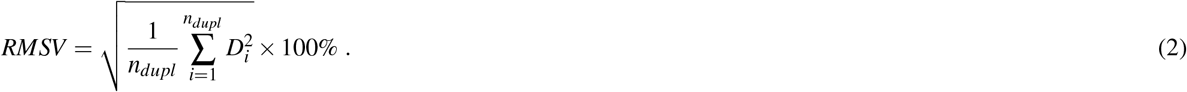

**Figure 2.**
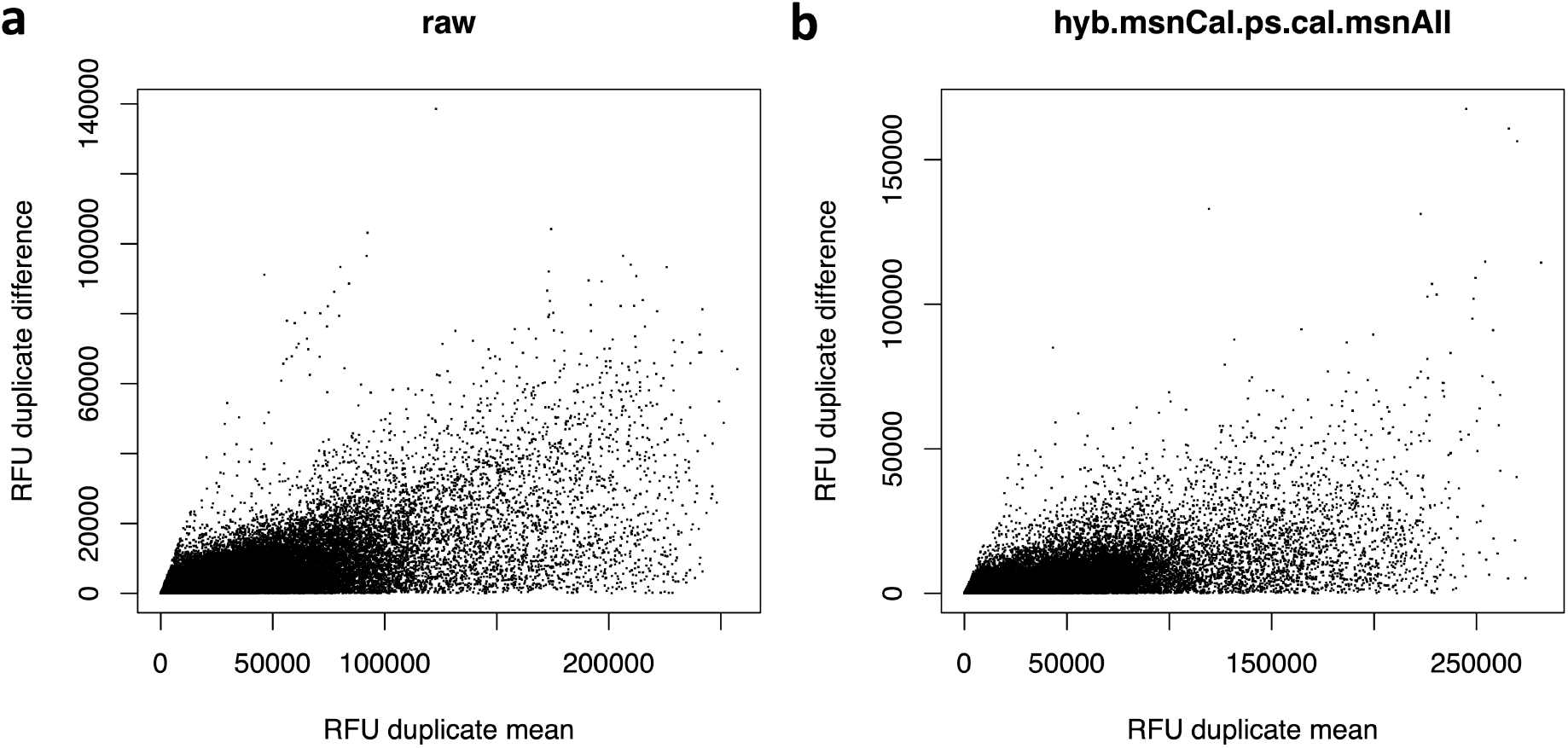
RFU spread increases with relative concentration. Bland-Altman plots showing the RFU difference (in absolute value) between inter-plate technical duplicates versus their corresponding mean. Each dot corresponds to the combination of one technical duplicate pair and one human protein SOMAmer, for a total of 102 duplicate pairs × 7,288 SOMAmers. **a** Raw data. **b** Fully normalized dataset (hyb.msnCal.ps.cal.msnAll).

By virtue of the sum of squares in the numerator, RMSV is sensitive to deviations from normality and to the presence of outliers. Alternatively, the Mean Absolute Difference Variation (MADV) metric is defined as

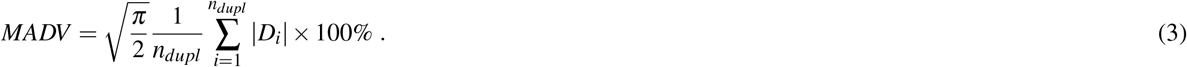

The 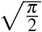 prefactor accounts for the relationship between MADV and the standard deviation of a Gaussian distribution. Since it scales linearly with the absolute value of the differences, this definition is less sensitive to outliers. A third definition, the Percentile Variation (PV) metric, is defined as

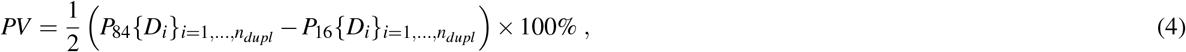

where *P*_84_ and *P*_16_ are the 84^th^ and 16^th^ percentiles in the distribution of scaled relative differences. Since, for a Gaussian distribution, 68% of the values are within one standard deviation from the mean, this approach ignores the distribution tails and is, therefore, the least sensitive to outliers. Notice that, if the scaled relative differences are normally distributed, these variability estimates are equivalent; the magnitude of the differences between these estimates is indicative of the magnitude and severity of deviations from normality.

Fig. 3 shows density distributions for each variability metric across all human protein SOMAmers, calculated from our *n*_*dupl*_ = 102 technical duplicates using different levels of normalization, as indicated. The first observation is that, regardless of metric, the sequential normalization approach appears to have a significant effect in succeeding to remove nuisance technical variability. Indeed, the median signal normalization procedure applied to all sample types leads to a fully normalized dataset (purple) that appears significantly less variable than datasets without it. Using median and MAD (*Median Absolute Deviation*) as robust statistics to characterize each distribution’s center and spread, we find that the variability of fully normalized datasets is 5.42 ± 2.13% for RMSV, 4.79 ± 1.80% for MADV, and 4.20 ± 1.56% for PV; hence, as anticipated by theoretical considerations, these distributions’ centers and widths are largest for RMSV and smallest for PV, with MADV in between those two. The variability metrics for each SOMAmer and normalization step are provided as Supplementary Data 2.

**Figure 3.**
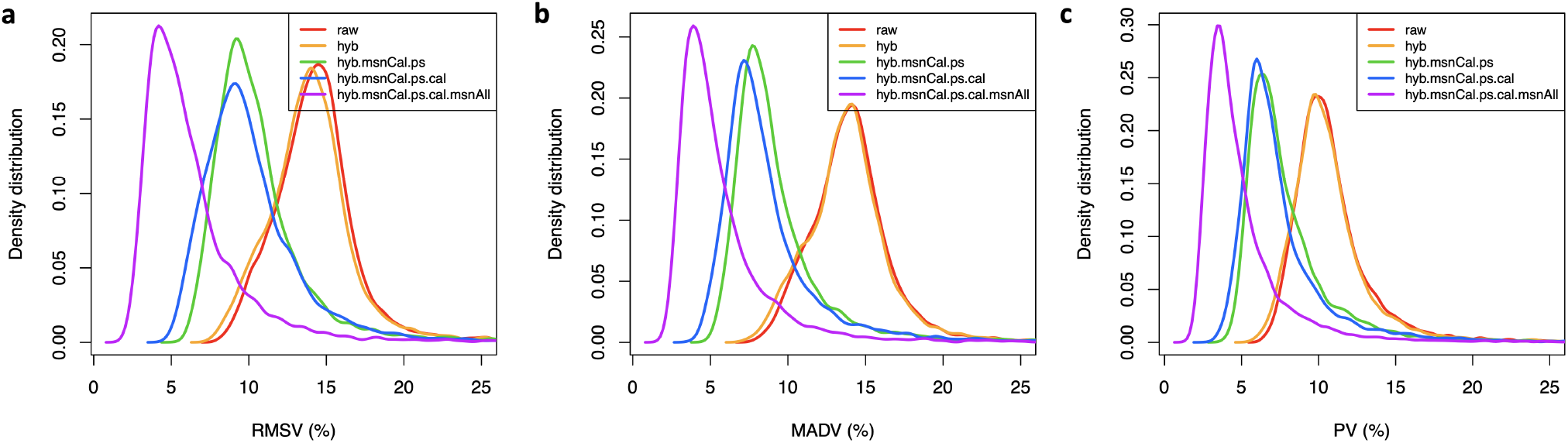
Assessment of SOMAmer variability. Density distributions representing different variability measurements across 7,288 human protein SOMAmers, obtained from inter-plate technical duplicates from 102 human participants. Each panel shows five distributions corresponding to different normalizations, as indicated. **a** Root-Mean-Squared Variation (RMSV). **b** Mean Absolute Difference Variation (MADV). **c** Percentile Variation (PV).

If we had many technical replicates from each participant at our disposal, a natural approach would be to characterize assay variability via the coefficient of variation (CV) defined as *CV* = (*σ*/*μ*) × 100%, i.e. the ratio between the standard deviation and the mean of all technical replicates. However, for technical duplicates, this definition is unstable because it relies on the calculation of standard deviations using just two data points per distribution (where each distribution describes replicate measurements from one subject-SOMAmer pair). Instead, here we propose a novel grid-search procedure that estimates CV from a large number (*n*_*dupl*_ = 102) of technical duplicates. To illustrate this approach, let us focus on Interleukin 8 (IL-8), also known as chemokine (C-X-C motif) ligand 8 (CXCL8), one of the major mediators of the inflammatory response, and Cystatin C, a protein associated with kidney function. These proteins have been extensively studied and constitute robust biomarkers for a variety of clinical conditions. Moreover, in the context of SomaScan, the SOMAmers for IL-8 and Cystatin C have been validated against antibody-based assays and established as useful biomarkers to predict clinical outcomes^14, 15, 18–20^.

The theoretical relationship between the coefficient of variation and the expected fold change (FC) ratio between repeated measurements was derived by Reed, Lynn, and Meade^21^. We assume that two replicate measurements, *RFU*_1_ and *RFU*_2_, are log-normally distributed random variables with the same mean and variance, and we define the fold change *FC ≡ RFU*_1_*/RFU*_2_ to be restricted to *FC* ≥ 1, without loss of generality. Then, the probability that these replicate measurements differ by a fold change equal or larger than *FC* is given by

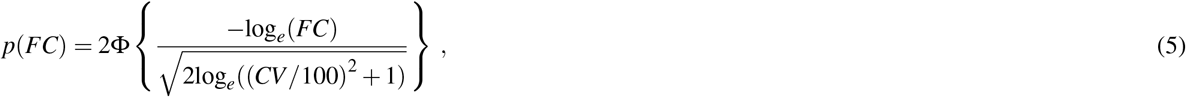

where Φ is the cumulative standard normal distribution function.

The top panels in Fig. 4 display theoretical estimates assuming different CVs (based on equation (5)), which are compared to distributions derived from our measured technical duplicates using different normalizations, as indicated. In agreement with our previous results, here we observe that the effect of subsequent normalization steps is that of reducing the fold-change differences between technical duplicates. In order to estimate the similarity between each one of these empirical distributions and each one of the theoretical distributions, we implement the non-parametric, one-sample Kolmogorov-Smirnov (KS) test. More specifically, we explore a dense ensemble of theoretical distributions generated by a pseudo-continuous scanning of the CV parameter in equation (5) and evaluate the similarity between each empirical distribution and each theoretical one as the *−log*_10_(p − value) derived from the one-sample KS test. For IL-8 and Cystatin C, this is shown by the bottom panels in Fig. 4. For each normalization, the estimated CV corresponds to the minimum of the corresponding curve, i.e. the CV that generates the theoretical distribution most similar to the measured one. As expected, these minima shift towards the left (i.e. smaller CV values) as we add normalization steps.

**Figure 4.**
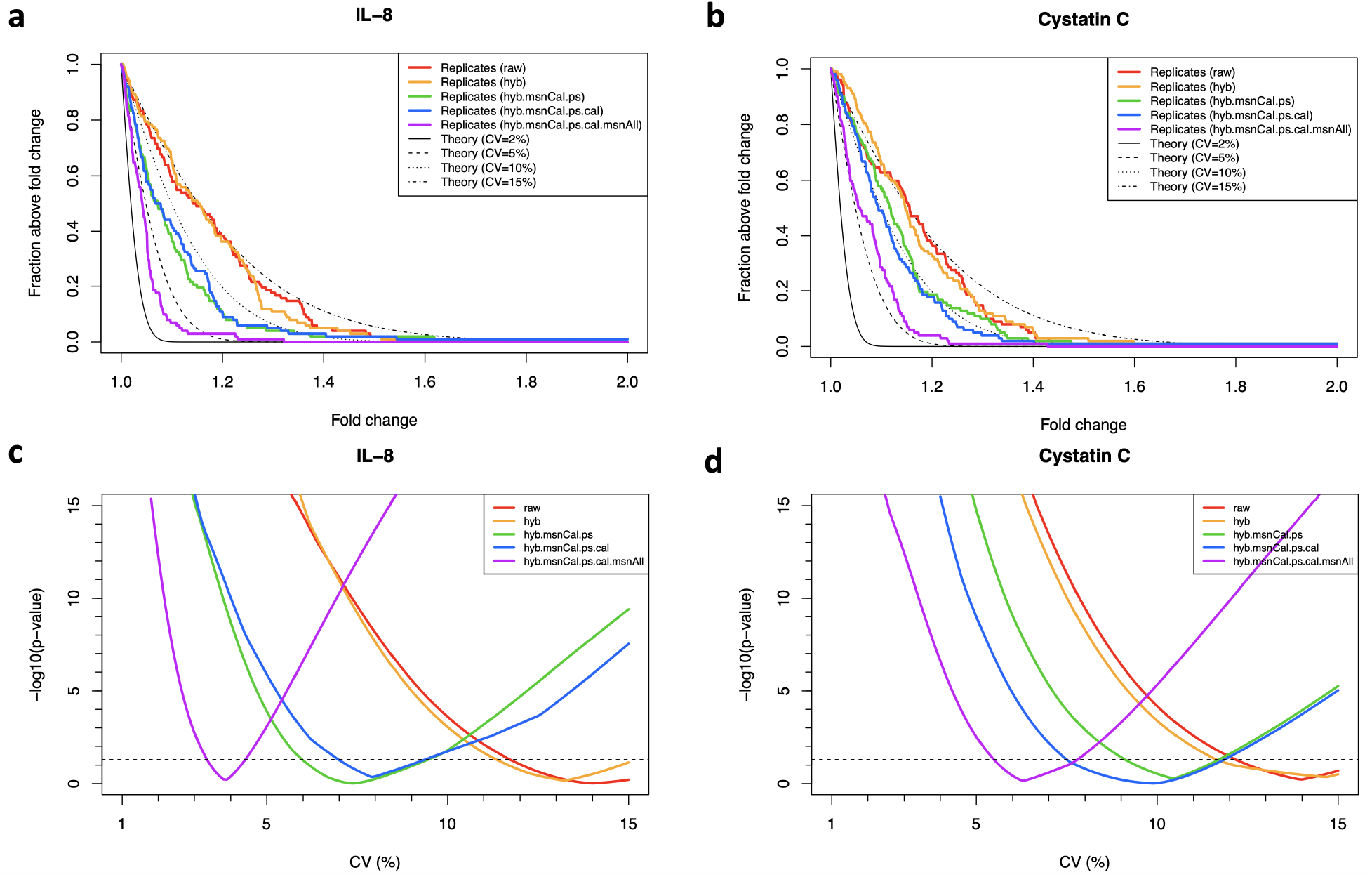
Grid-search procedure to determine the coefficient of variation (CV) from inter-plate technical duplicates. The method is here illustrated with IL-8 **(a,c)** and Cystatin C **(b,d)**. **a-b** Probability that two replicate measurements differ by a factor larger than a given fold change. Theoretical estimates assuming different CVs (based on equation (5)) are compared to technical duplicates obtained from 102 human subjects using different normalizations, as indicated. **c-d***−log*_10_(p − value) from Kolmogorov-Smirnov tests comparing theoretical and measured distributions as a function of the assumed CV. The minimum in each curve indicates the CV that yields the theoretical distribution most similar to the measured one. Horizontal dashed lines correspond to p − value = 0.05.

By applying this grid-search procedure to all human protein SOMAmers, we obtain the CV density distributions for all normalizations shown in Fig. 5(a). In more detail, Fig. 5 (b-f) display cumulative distributions of the number of SOMAmers below a given CV for each normalization. Here, we observe that the trends illustrated by IL-8 and Cystatin C are indeed confirmed across all SOMAmers in the assay, and are also consistent with our previous analysis based on other variability metrics (Fig. 3). The estimated CVs for each SOMAmer and normalization step are provided as Supplementary Data 3.

**Figure 5.**
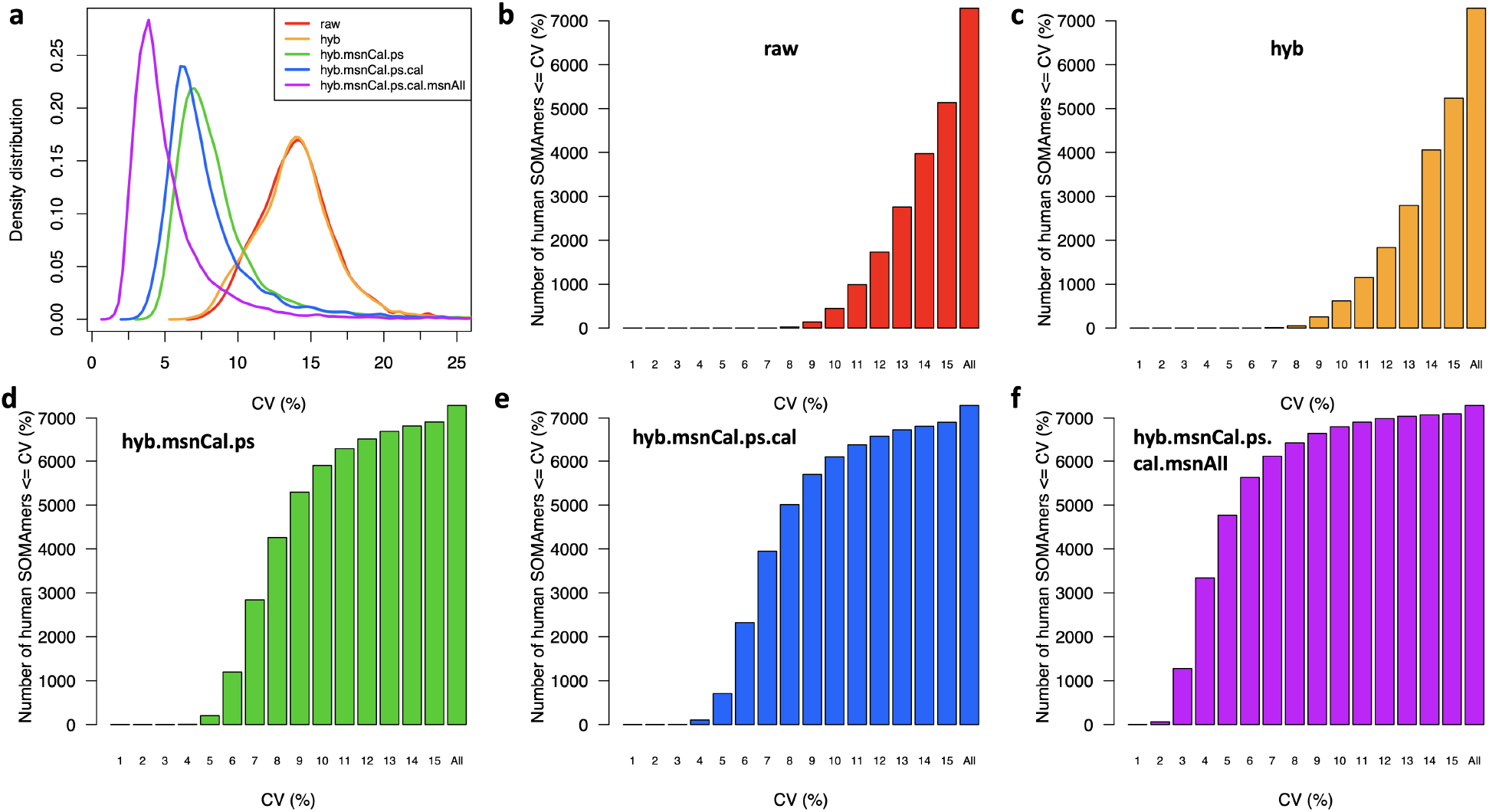
Distributions of coefficients of variation (CV). **a** Density distributions of CV across 7,288 human protein SOMAmers, obtained from inter-plate technical duplicates from 102 human subjects for different normalizations. **b-f** Cumulative distribution of the number of SOMAmers below a given CV for different normalizations.

Fig. 6 shows the CV as a function of the median signal-to-background ratio (SBR, defined as the median RFU of all experimental samples divided by the median RFU of all buffer wells) for all human protein SOMAmers, colored by dilution group. There is a significant correlation between CV and SBR (Spearman’s *r* = 0.35, *p* = 4 × 10^−210^) primarily driven by the 20% dilution group (*r* = 0.43, *p* = 2 × 10^−267^), which represents 82% of all human protein SOMAmers, followed by the 0.5% dilution group (*r* = 0.14, *p* = 4 × 10^−6^), which represents 15% of all human protein SOMAmers. The third group, 0.005% dilution, comprises only 3% of all human protein SOMAmers and shows no significant correlation between CV and SBR. The CV median ± MAD across SOMAmers are 4.5 ± 1.5% (for all human protein SOMAmers), 4.5 ± 1.5% (for the 20% dilution group), 4.0 ± 1.5% (for the 0.5% dilution group), and 6.5 ± 2.2% (for the 0.005% dilution group), respectively. The SBR median ± MAD across SOMAmers are 8.1 ± 7.2 (for all human protein SOMAmers), 6.7 ± 5.1 (for the 20% dilution group), 26.6 ± 26.7 (for the 0.5% dilution group), and 76.2 ± 96.2 (for the 0.005% dilution group), respectively. Pairwise Wilcoxon test comparisons between dilution groups indicate that all these CV and SBR differences are statistically significant (adjusted p-value < 0.05). Notice that the data shown in Fig. 6 correspond to the full normalization (hyb.msnCal.ps.cal). Results for all SOMAmers and normalization steps are provided as Supplementary Data 4.

**Figure 6.**
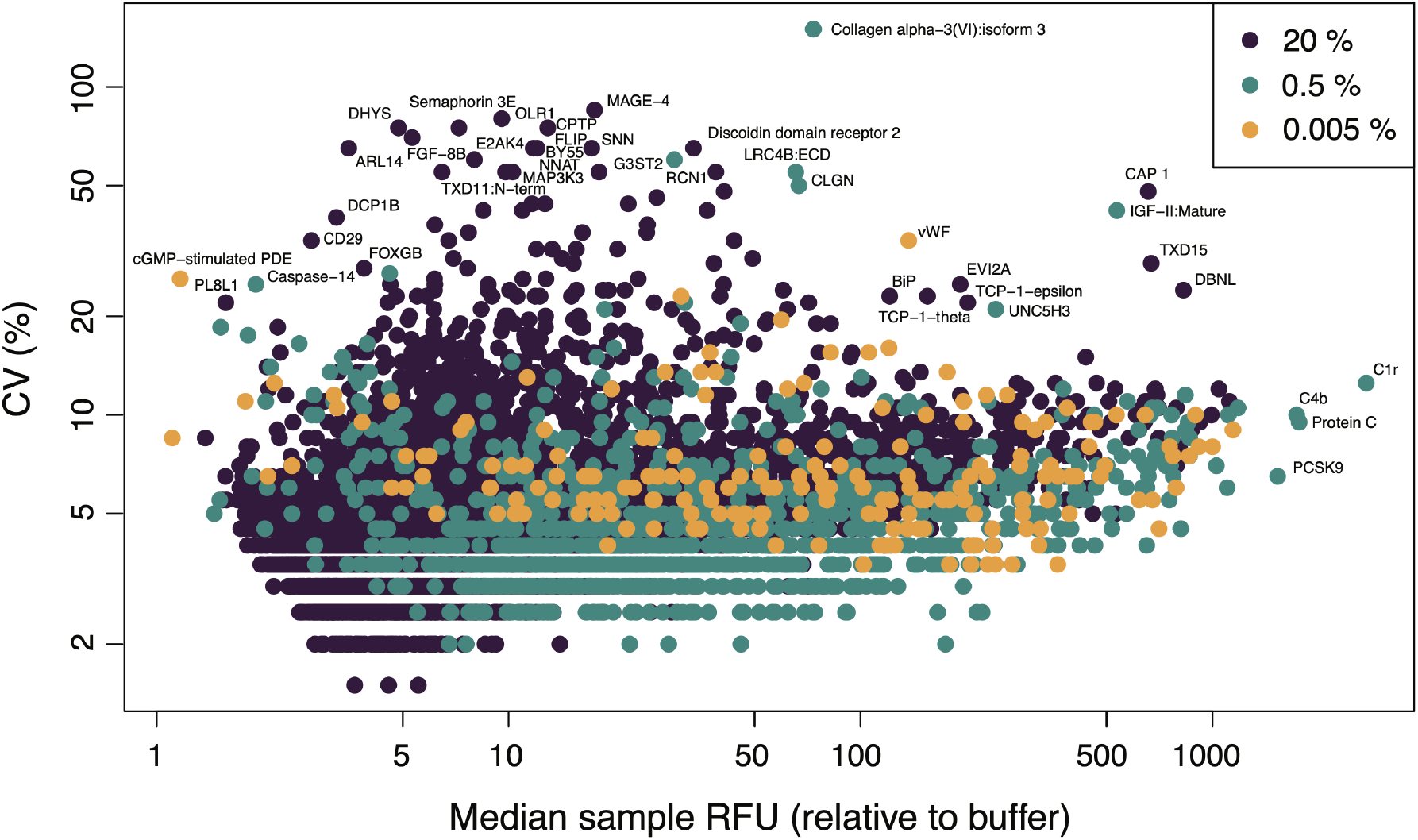
Brightness and variability of SOMAmers by dilution group. Coefficient of variation of 7,288 human protein SOMAmers as a function of the median signal-to-background ratio across experimental samples. SOMAmers are colored based on dilution: 20% (*n* = 6001), 0.5% (*n* = 1100), and 0.005% (*n* = 187). Data have been fully normalized.

A remarkable characteristic of SomaScan is its sensitivity: without reporting any missing values (which constitutes, in its own right, a major departure from all antibody-based and most other proteomic assays), relative concentrations measured in experimental samples turn out to be significantly higher than background for most SOMAmers. Indeed, 99.1% of human protein SOMAmers are > 2−fold brighter in experimental samples compared to buffer wells, 68.8% of them are 5−fold brighter, and 43.8% of them are even 10−fold brighter and beyond. It should be pointed out, however, that RFU values below the limit of detection (LoD) should effectively be disregarded. Derived by Armbruster and Pry^22^ based on the Clinical and Laboratory Standards Institute definition, the estimated limit of detection (eLoD) for the *α*-th SOMAmer can be calculated via robust statistics as

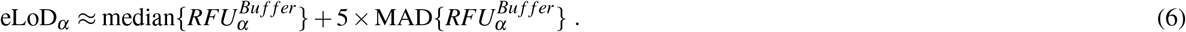

In our dataset, we find that the median RFU of experimental samples is below eLoD for only seven human protein SOMAmers, whose targets are Caspase-14, PL8L1, ERGI1, CYBR1, DC-SIGNR, cGMP-stimulated PDE, and U773 (see Supplementary Data 4 for more details).

Despite the remarkable sensitivity, however, more scrutiny has been placed on SomaScan’s specificity, cross-reactivity, and orthogonal assay reproducibility (see Discussion section below and references therein). Although a full investigation of these issues lies beyond the scope of the present paper, our study provides some insight from the analysis of SOMAmer controls. Fig. 7 shows the RFU median and 95% CI range for experimental samples (blue) and buffer wells (black) for various types of control SOMAmer available in the assay, as well as the corresponding eLoD (red) estimated via equation (6). As described in the Methods section, Hybridization Control Elution SOMAmers (*n* = 12) are included in each well to adjust for noise in the hybridization process. Thus, the same signal is expected from buffer wells and experimental samples, which is confirmed in our data. Other control SOMAmers not expected to generate signals above buffer are described as non-cleavable (*n* = 4), non-biotin (*n* = 10), and spuriomers (*n* = 20), purposefully designed to interfere with different phases of the experimental process. Surprisingly, however, we observe excess signals for these control SOMAmers in a significant fraction of experimental samples compared with those from buffer wells. In terms of the median signal-to-background ratio (SBR) defined above, we find that, while all non-cleavable controls and most spuriomers show SBR < 2, some spuriomers lie in the SBR = 2 − 3 range, and more pronounced excess signals are detected from non-biotin SOMAmers.

**Figure 7.**
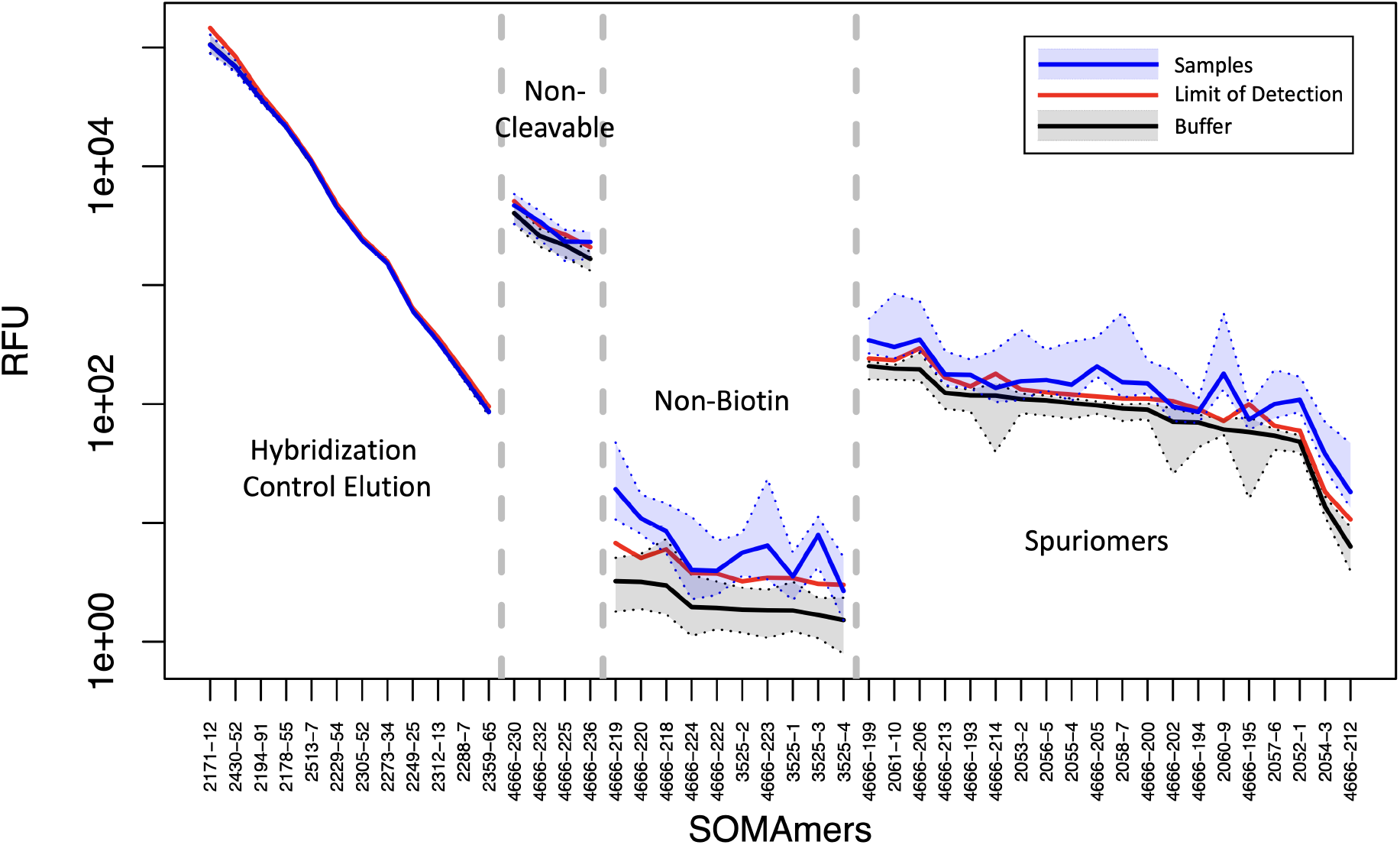
Signal-to-background assessment of control SOMAmers. Intensity (measured in Relative Fluorescence Units) for different types of control SOMAmer, as indicated. The median (solid line) and 95% CI (shaded area) are shown for buffer wells (black) and experimental samples (blue). The estimated Limit of Detection derived from equation (6) is shown as a solid red line. Control SOMAmers within each type are shown in decreasing order of median RFU intensity in buffer wells. Data have been fully normalized.

SOMAmer reagents bind cognate proteins in their native folded conformations and may, therefore, be sensitive to the complex variety of proteoforms arising from protein-altering DNA variants, from alternative splicing of RNA transcripts, and from many types of post-translational modification. Most, but not all, SOMAmers are in a one-to-one relationship with their annotated cognate proteins (see Methods section for details). In cases where two or more different SOMAmers share the same annotated target, we calculated the pairwise RFU correlation across experimental samples in our cohort (Supplementary Fig. 2, Supplementary Data 5). We observe two distinct correlation distributions, namely, one of them approximately centered around *r* ≈ 0, and another one that peaks at *r* ≈ 1. The latter indicates the expected redundancy of SOMAmers that bind to the same target, whereas the former may point to SOMAmers that appear sensitive to proteoform complexity. This important topic certainly deserves further investigation that lies beyond the scope of our present work. The analyses presented here are based on individual SOMAmers and are not dependent upon how targets are annotated.

## Discussion

Based on the analysis of more than 2,000 samples in 22 plates, we presented an extensive assessment of the current plasma 7k SomaScan proteomics assay. In the first place, we outlined a sequence of normalization steps that account for different sources of nuisance variance. Although similar to analysis pipelines described previously^8, 12^ and to those implemented by SomaLogic’s informatics, the updated pipeline described here is entirely self-contained and reproducible, independent from external references. For our cohort, the fully normalized datasets obtained with our pipeline are consistent with those provided by SomaLogic. The ability to run independent normalization analyses, however, is important for various reasons. On the one hand, the set of external references (based on a fixed pool of healthy human control samples) may be inappropriate to study individuals and populations with large deviations from a healthy plasma proteome. As noted by Lopez-Silva et al^20^ in their study of chronic kidney disease, SomaLogic’s normalization could potentially attenuate the strength of associations by reducing more extreme values if they are associated with a clinical outcome, perhaps contributing to the attenuated prognostic associations they observed with SomaScan. Pietzner et al^16^ report that using SomaScan data without a normalization step applied to correct for unwanted technical variation and to make data comparable across cohorts, a higher median correlation with results from the Olink proteomics assay^23^ was observed, along with substantial differences in the association with various phenotypic characteristics. On the other hand, users may be interested in using their own set of control samples to bridge across studies, thereby needing to calibrate different studies using their own controls. More generally, users may be interested in adapting the normalization process according to their studies’ characteristics and objectives. Therefore, we believe that presenting this independent normalization pipeline empowers SomaScan end users to tailor the normalization procedures to their own individual needs.

Secondly, we assessed different normalization approaches with an independent set of inter-plate technical duplicates from 102 donors. As opposed to using pooled plasma, which has the effect of averaging out naturally occurring proteomic differences across individual subjects, our study was designed to capture those differences. A variety of approaches (PCA, different variation metrics, and a novel framework to estimate CV based on a large number of technical duplicates) confirmed that median signal normalization, applied on all sample types *after* removing nuisance inter-plate effects via calibrators, appears indeed as the optimal approach to minimize technical variability. The overall CV median ± MAD across all human protein SOMAmers is 4.5 ± 1.5%.

Thirdly, we analyzed signal-to-background ratios, highlighting the remarkable sensitivity of SomaScan. More than 99% of human proteins are > 2−fold brighter in experimental samples compared to buffer wells. Different from other proteomic platforms, SomaLogic’s informatics does not provide limit of detection assessments; all SOMAmer/sample combinations are quantified with an RFU measurement. By including an estimated limit of detection (eLoD) based on buffer wells, we found only 7 human protein SOMAmers in our samples whose median RFU lied below eLoD.

Next, we analyzed various types of control SOMAmers included in the assay. For many of these control SOMAmers, we found that the signal from our samples was higher than that measured in buffer wells. This surprising finding was most noticeable among so-called non-biotin and spuriomers. Some of these spurious signals arose for low RFU values (ranging in the tens and hundreds); it is therefore plausible that some of those low-range RFU measurements are spurious and, thus, reporting numerical RFU values for all sample/SOMAmer combinations is misleading due to under-estimated experimental noise. In our study, RFU values adjusted by dilution over 1,799 experimental samples and 7,288 human protein SOMAmers span a fold-change range of 1.2 × 10^9^ (for the fully normalized dataset), which represents the dynamic range of measured concentrations. However, if low RFU values are unreliable due to background noise, the effective dynamic range should shrink by some additional 1-2 orders of magnitude in protein concentration. SomaLogic’s claim stating that the SomaScan assay is capable of quantifying relative protein abundances across 10 orders of magnitude remains, therefore, open to further examination.

As mentioned above, multiple studies have investigated SomaScan’s specificity, cross-reactivity, and orthogonal assay reproducibility, most of them performed on previous versions of the SomaScan assay. Emilsson et al.^4^ ran a Novartis custom-designed 5k serum SomaScan assay on 5,457 participants from the AGES Reykjavik study and performed a direct validation using two different mass spectrometry techniques, namely data dependent analysis (DDA) and multiple reaction monitoring (MRM). Results of the mass spectrometry experiments provided confirmatory evidence of 779 SOMAmer reagents binding their endogenous targets (736 by DDA and 104 by MRM). Sun et al.^5^ performed 4k plasma SomaScan on 3,301 healthy participants from the INTERVAL study. They identified 1,927 genotype–protein associations (pQTLs), and tested 163 of them with Olink, finding strongly correlated effect-size estimates (*r* = 0.83). To assess potential off-target cross-reactivity, they also tested 920 SOMAmers for detection of proteins with at least 40% sequence homology to the target protein. Although 126 (14%) SOMAmers showed comparable binding with a homologous protein, nearly half of these were binding to alternative forms of the same protein. Graumann et al.^24^ compared 1.3k plasma SomaScan against 368 proteins from Olink in 20 high-grade serous carcinoma patients and 20 individuals with non-malignant gynecologic disorders, followed by validation with ELISA. Raffield et al.^15^ analyzed 1.1k and 1.3k serum and plasma SomaScan in comparison with Myriad RBM Luminex, Meso Scale Discovery, Olink, and ProterixBio assays in multiple cohorts (SPIROMICS, COPDGene, MESA, and others). Pietzner et al.^16^ performed 5k plasma SomaScan and compared it with 1069 protein targets from 12 Olink panels in 485 participants in the Fenland study. Lim et al.^18^ performed 1.3k plasma SomaScan compared with the 65-plex bead-based Discovery (Eve Technologies) immuno-assay in 24 melanoma patients treated with combination immune checkpoint inhibitors, followed by validation with the Milliplex bead-based immuno-assay. Kukova et al.^19^ performed 1.3k plasma and urine SomaScan compared with various immuno-assays (Randox Biochip Array, Abbott Architect, Meso Scale Discovery, UniCel) in pre- and post-operative samples from 54 individuals with acute kidney injury after cardiac surgery. Tin et al.^14^ performed 4k plasma SomaScan compared with 9 proteins measured via clinical assays in 42 participants in the ARIC longitudinal study. Very recently, Lopez-Silva et al.^20^ performed 7k serum SomaScan compared with 9 proteins measured by immuno-assays in 498 participants in the African American Study of Kidney Disease and Hypertension (AASK). Although not intending to exhaustively review the SomaScan literature, these references depict the wide range of approaches pursued to characterize the performance of the SomaScan’s assay and to link it to established clinical chemistry assays, as well as to other experimental platforms.

Let us discuss the limitations of this study. As stated above, our assessment relies on a large number of samples across multiple plates, which, to the best of our knowledge, is the largest plasma 7k SomaScan study published to date. Variability was estimated using inter-plate technical duplicates from 102 human donors. A full intra- and inter-plate assessment requires the use of multiple technical replicates, typically three or more, within and across plates, which were not available within the scope of this study. A more exhaustive characterization, moreover, would require the use of a sequence of serial dilutions of a standard sample within each plate to evaluate the linearity of the measured signal versus protein concentration, as well as the limit of detection and limit of quantitation. Such procedure, which is standard in other proteomic assays e.g. immuno-assays, would be prohibitively costly in SomaScan. On the other hand, spike-in experiments^25, 26^ could be carried out to assess the assay’s ability to recapitulate predefined concentration differences among sample groups. The setup described in this work, however, follows SomaLogic’s plate design recommendations and reflects the typical setup of a SomaScan experiment. As described above, previous studies using earlier versions of the SomaScan technology investigated other technical aspects that lie beyond the scope of our present work. These involve orthogonal validations using other proteomic assays such as mass spectrometry, immuno-assays and Olink, as well as off-target cross-reactivity assessments.

Certainly, more work is needed to fully address background noise, limits of detection, specificity, cross-reactivity, and orthogonal reproducibility. We conclude here, nevertheless, by emphasizing the tremendously promising opportunities offered by the SomaScan assay, with its increasingly expanding protein coverage and consistently low variability. These opportunities include (i) biomarker discovery and the elucidation of biological pathways in response to disease or treatment, (ii) uncovering the complex connections between genome and proteome, and (iii) the assay’s potential as a companion diagnostic and tool for patient stratification^2^. We hope this innovative platform will continue to improve and grow, thereby aiding basic, translational, and clinical research communities in their quest to understand human health and conquer disease.

## Methods

### Experimental

A total of 1,806 human plasma samples were obtained from participants in the Baltimore Longitudinal Study of Aging (BLSA), which is supported by the National Institute on Aging (NIA) at the NIH. Proteomic profiles were characterized using the 7k SomaScan assay v4.1 (SomaLogic, Inc.; Boulder, CO, USA). This assay consists of 7,596 SOMAmers, out of which 7,288 target annotated human proteins. The remaining SOMAmers are different types of control, including 12 HCE (*Hybridization Control Elution*) SOMAmers used in the Hybridization control normalization step (described below), 4 non-cleavable SOMAmers, 10 non-biotin SOMAmers, 20 spuriomers (designed as random, non-specific sequence motifs), and multiple legacy SOMAmers targeting non-human proteins. In order to cover a broad range of endogenous concentrations, SOMAmers are binned into different dilution groups, namely 20% (1:5) dilution for proteins typically observed in the femto- to pico-molar range (which comprise 82% of all human protein SOMAmers in the assay), 0.5% (1:200) dilution for proteins typically present in nano-molar concentrations (15% of human protein SOMAmers in the assay), and 0.005% (1:20,000) dilution for proteins in micro-molar concentrations (3% of human protein SOMAmers in the assay). The human plasma or serum volume required is 55 *μ*L per sample.

SOMAmers are uniquely identified by their “SeqId”, but the relation between SOMAmers and annotated proteins is not one-to-one. Indeed, the 7,288 SOMAmers that target human proteins are mapped to 6,383 unique UniProt IDs, 6,373 unique Entrez gene IDs and 6,365 unique Entrez gene symbols. In addition, 233 SOMAmers target mouse proteins. All of the SOMAmers in the 5k SomaScan assay v4 (as well as most of those in the 1.3k SomaScan assay v3) are included in the 7k assay (Supplementary Figure 1). Data processing and delivery by SomaLogic is in the adat format^27^, and it remains consistent across assay versions. The data normalization procedures described below are equally applicable to all assay versions. It should be pointed out, however, that the 1.3k assay has a different dilution scheme, with 40%, 1%, and 0.005% dilution groups. For more details, see Supplementary Data 1.

The experimental procedure follows a sequence of steps, namely: (1) SOMAmers are synthesized with a fluorophore, photocleavable linker, and biotin; (2) diluted samples are incubated with dilution-specific SOMAmers bound to streptavidin beads; (3) unbound proteins are washed away, and bound proteins are tagged with biotin; (4) UV light breaks the photocleavable linker, releasing complexes back into solution; (5) non-specific complexes dissociate while specific complexes remain bound; (6) a polyanionic competitor is added to prevent rebinding of non-specific complexes; (7) biotinylated proteins (and bound SOMAmers) are captured on new streptavidin beads; and (8) after SOMAmers are released from the complexes by denaturing the proteins, fluorophores are measured following hybridization to complementary sequences on a microarray chip. The fluorescence intensity detected on the microarray, measured in RFU (*Relative Fluorescence Units*), is assumed to reflect the amount of available epitope in the original sample. This workflow is summarized in Supplementary Fig. 3.

Samples are organized in 96−well plates, each plate consisting of buffer wells (with no sample added), calibrator and QC samples (provided by SomaLogic from pooled healthy donor controls), and the experimental samples of interest. In this study, a total of 2,050 samples (corresponding to 68 buffer wells, 110 calibrators, 66 QC, and 1,806 human plasma samples from BLSA participants) were analyzed across 22 plates. Based on observed issues during the experiment (low sample volume, clogged vial, etc) as well as on PCA (*Principal Component Analysis*) results, 7 outlier experimental samples were removed from the downstream analysis. Of the remaining 1,799 experimental samples, 204 corresponded to inter-plate technical duplicates from 102 BLSA participants.

### Data Normalization Procedures

Raw data, as obtained after aggregation from slide-based hybridization microarrays, exhibit intra-plate nuisance variance due to differences in loading volume, leaks, washing conditions, etc, which is then compounded with batch effects across plates. In order to account for intra- and inter-plate variability of buffer, calibrator, QC, and experimental samples, here we consider a sequence of steps, whose main elements are hybridization normalization, median signal normalization, plate-scale normalization, and inter-plate calibration. Each of these steps generates scale factors at different levels: plate-specific, by SOMAmer dilution group, SOMAmer-specific, by sample type, and combinations thereof. Besides removing technical variability, these scale factors can be used as quality control flags at the plate-, sample-, and SOMAmer-levels^8^. A summary of these data normalization procedures, explained below in detail, is shown in Table 1. To keep a consistent mathematical notation in the expressions below, wells are indicated by Latin sub-indices and SOMAmers by Greek sub-indices. Plate index, sample type, and SOMAmer groupings are denoted by super-indices. We use *well* and *sample* interchangeably, although it should be noticed that no sample material is added to buffer wells; also, depending on context, *sample* may refer specifically to the experimental samples of interest, thus excluding control sample types such as buffer, calibrator, and QC.

**Table 1.**
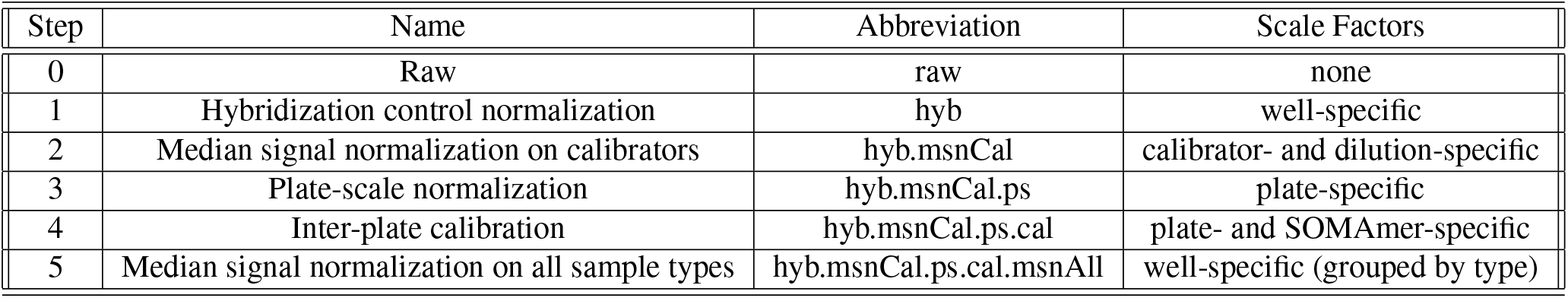
Summary of normalization steps.

#### 1. Hybridization control normalization (hyb)

Hybridization control normalization is designed to adjust for nuisance variance on the basis of individual wells. Each well contains *n*_*HCE*_ = 12 HCE (*Hybridization Control Elution*) SOMAmers at different concentrations spanning more than 3 orders of magnitude. By comparing each observed HCE probe to its corresponding reference value, and then calculating the median over all HCE probes, we obtain the scale factor for the *i*-th well, i.e.

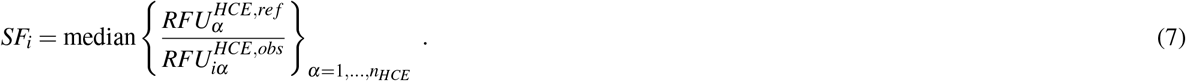

Notice that this normalization step is performed independently for each well; once the scale factor is determined, all SOMAmer RFUs in the well are multiplied by the same scale factor. Instead of using an external reference, a plate-specific internal reference can be determined by the median across the *n*_*s*_ = 96 wells in the plate, i.e.

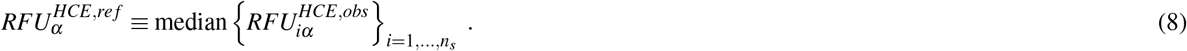

#### 2. Median signal normalization on calibrators (hyb.msnCal)

Median signal normalization is an intra-plate normalization procedure performed within wells of the same sample class (i.e. separately for buffer, QC, calibrator, and experimental samples) and within SOMAmers of the same dilution grouping (i.e. 20%, 0.5%, and 0.005%). It is intended to remove sample-to-sample differences in total RFU brightness that may be due to differences in overall protein concentration, pipetting variation, variation in reagent concentrations, assay timing, and other sources of variability within a group of otherwise comparable samples. Since RFU brightness differs significantly across SOMAmers, median signal normalization proceeds in two steps. First, the median RFU of each SOMAmer is determined (across all samples of the same sample type) and sample RFUs are divided by it. The ratio corresponding to the *i*-th sample and *α*-th SOMAmer is thus given by

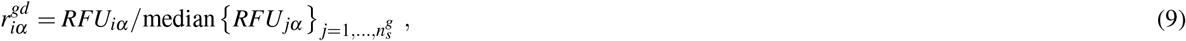

where indices *g* and *d* denote sample type and SOMAmer dilution groupings, respectively. Then, the scale factor associated with the *i*-th sample is determined as the inverse of the median ratio for that sample across all SOMAmers in the dilution group:

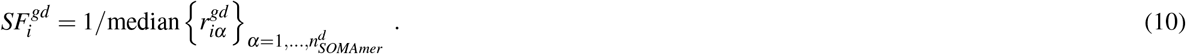

To median-normalize the *i*-th sample, then, all its SOMAmer RFUs in the same dilution group are multiplied by this scale factor. As discussed in our previous work^12^, performing median signal normalization on experimental samples *before* inter-plate calibration presents the risk of enhancing plate-to-plate differences. Thus, in this step, we restrict median signal normalization to calibrators only.

#### 3. Plate-scale normalization (hyb.msnCal.ps)

Plate-scale normalization aims to control for variance in total signal intensity from plate to plate. No protein spikes are added to the calibrator; the procedure solely relies on the endogenous levels of each protein within the set of calibrator replicates.

For the *α*-th SOMAmer on the *p*-th plate,

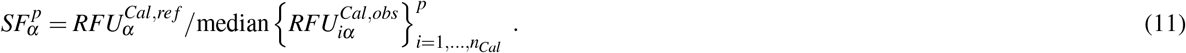

Calibration scale factors may be pinned to an external reference, but here we utilize an internal reference determined by the median across all calibrators on all *n*_*p*_ plates, i.e.

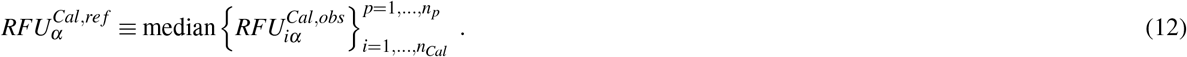

In order to correct the overall brightness level of the *p*-th plate, we calculate the plate-scale scale factor as the median of 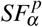 across all SOMAmers, i.e.

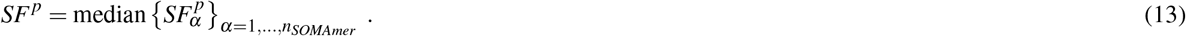

For all wells on the *p*-th plate and all SOMAmers, RFUs are multiplied by the plate-scale factor *SF^p^*.

#### 4. Inter-plate calibration (hyb.msnCal.ps.cal)

Following plate-scale normalization, we recalculate SOMAmer- and plate-specific scale factors via Eqs. (11)-(12). Separately for each SOMAmer and plate, all wells on the plate are corrected by the recalculated 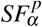.

#### 5. Median signal normalization on all sample types (hyb.msnCal.ps.cal.msnAll)

At this stage, after correcting plate-to-plate variability to the fullest extent possible, median signal normalization (described in step 2) can be performed separately on each sample type. This step yields the final, fully-normalized dataset. It should be pointed out that SomaLogic currently delivers normalized output files (as plain text files of extension .adat^27^) that follow similar steps as those described here, but using external references^8^. These references may be outdated (because they are specific to the control samples used), not necessarily representative of the target samples of interest (because they utilize a fixed pool of healthy human control samples for the last normalization step) and are not delivered with the .adat files provided to the customer (therefore precluding any attempt of independent analysis). In contrast, the process described here relies solely on internal references derived from the measured data. For the data analyzed in this study, fully normalized datasets using internal versus external references are highly concordant; for all human protein SOMAmers in the assay, Spearman’s RFU correlation across all experimental samples yields *r* ≥ 0.95 and *p* ≈ 0. It should be noted that, due to Spearman’s correlation invariance, these results remain valid under any type of monotonic (e.g. logarithmic) RFU transformations.

## Computational

Analyses were performed in R version 4.1.2. Packages used were: RColorBrewer (v. 1.1.2), viridis (v. 0.6.2), MASS (v. 7.3.54), calibrate (v. 1.7.7), VennDiagram (v. 1.7.3).

## Statistical

Unless explicitly noted otherwise, statistical tests were two-sided.

## Data Availability

Anonymized datasets used for this technical study will be available (upon acceptance of this paper for publication) on the Open Science Framework repository, osf.io/[TBD]. Results from our analyses are provided as Supplementary Data files.

## Ethical approval and informed consent

The study protocol for this study was conducted in accordance with Declaration of Helsinki principles and was reviewed and approved by the Institutional Review Board of the National Institutes of Health’s Intramural Research Program. Written informed consent was obtained from all participants.

## Acknowledgements

This research was supported entirely by the Intramural Research Program of the National Institute on Aging, NIH. We thank Brian Sellers (NIH), Kathryn Jenko and Jennifer Yoon (SomaLogic, Inc.) for useful discussions.

## Author contributions

J.C. has designed the analysis, performed the analysis, and written the paper. G.N.D., T.T. have contributed to the conceptualization and methodology. L.F., K.A.W. have designed the study, provided resources, and acquired funding. All authors have reviewed the manuscript.

