## Supplementary Figures for "Assessment of Variability in the Plasma 7k SomaScan Proteomics Assay"

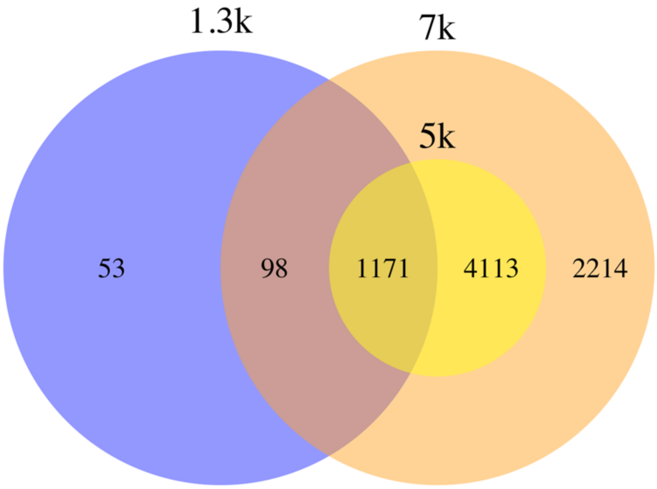


**Supplementary Figure 1.** Venn diagram showing the SOMAmer overlap (based on the unique "SeqId" identifiers) between the 1.3k (v3), 5k (v4), and 7k (v4.1) SomaScan assays. For more details, see **Supplementary Data 1**.

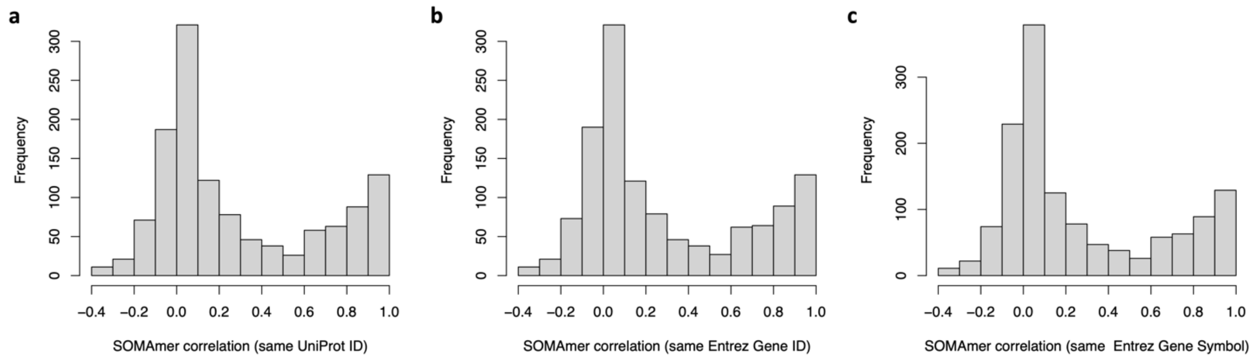


**Supplementary Figure 2.** Frequency distributions of pairwise Pearson's correlation of $log_{10}$(RFU) across 1,799 experimental samples between SOMAmers that share the same annotated target proteins according to: **(a)** UniProt ID, **(b)** Entrez Gene ID, and **(c)** Entrez Gene Symbol. For more details, see **Supplementary Data 5**.

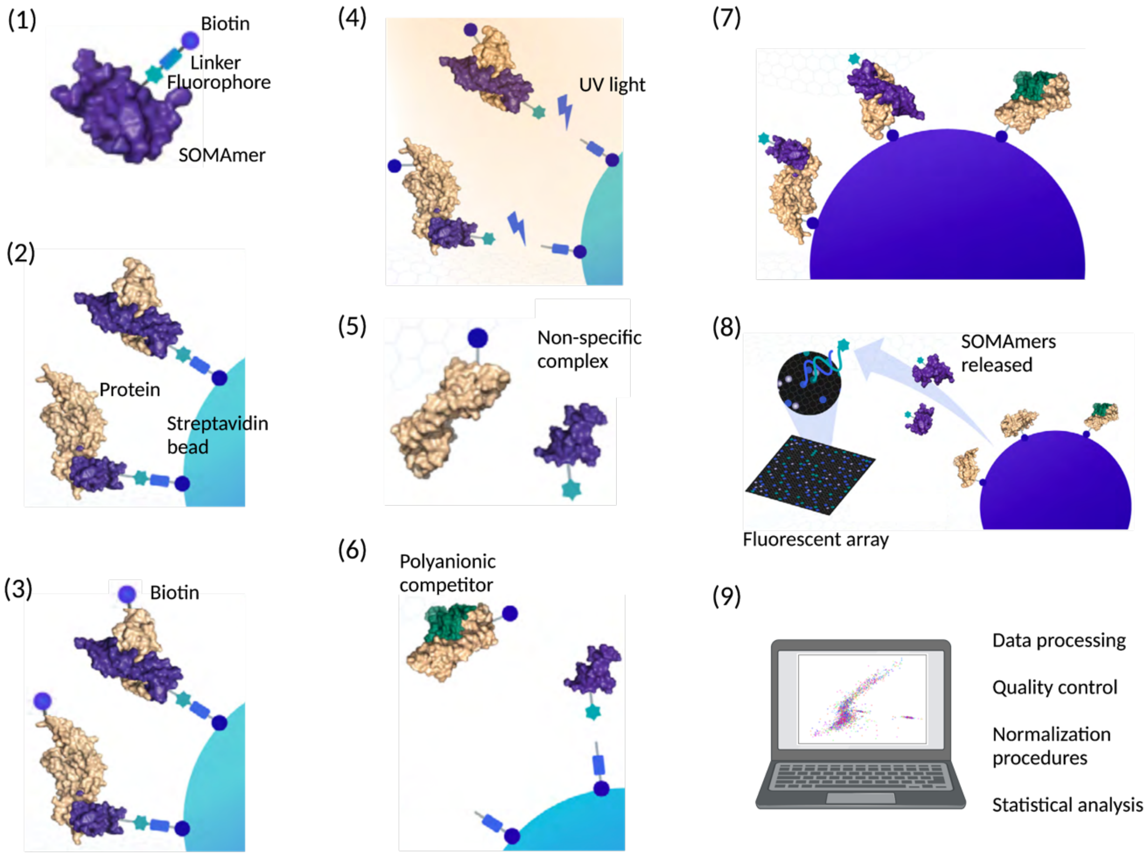


**Supplementary Figure 3.** Workflow of the assay. (1) SOMAmers are synthesized with a fluorophore, photocleavable linker, and biotin; (2) diluted samples are incubated with dilution-specific SOMAmers bound to streptavidin beads; (3) unbound proteins are washed away, and bound proteins are tagged with biotin; (4) UV light breaks the photocleavable linker, releasing complexes back into solution; (5) non-specific complexes dissociate while specific complexes remain bound; (6) a polyanionic competitor is added to prevent rebinding of non-specific complexes; (7) biotinylated proteins (and bound SOMAmers) are captured on new streptavidin beads; and (8) after SOMAmers are released from the complexes by denaturing the proteins, fluorophores are measured following hybridization to complementary sequences on a microarray chip. Upon completion of all experimental steps, (9) the bioinformatic analysis proceeds. Adapted from SomaLogic's Technical Note SL00000572 and created with BioRender.
